# Experimental evaluation of Eastern box turtle (*Terrapene carolina carolina*) detectability in visual search surveys

**DOI:** 10.1101/2024.01.02.572695

**Authors:** William Heinle, Noelle Beswick, Emily Wapman, J. Andrew Royle

## Abstract

Understanding how detection probability varies over time, space, or in response to measurable covariates is important to inform the monitoring and assessment of many species. A standard model to understand detectability – the availability/perception model – admits that detection probability is the composite of two components: availability and perception. Availability is largely affected by environmental and behavioral factors, whereas perception is primarily affected by attributes of individual observers and survey protocols, and thus can potentially be partially controlled by survey design. We designed and implemented a field study to understand the perception component of detection for Eastern box turtles (*Terrapene carolina carolina*) using visual encounter surveys. We obtained and deployed museum specimens of Eastern box turtle shells and subjected them to visual search surveys by observers in realistic field situations. Overall, about 50% of the box turtle shells were detected by observers including, 41.5% in the ‘partially visible’ state and 63% in the ‘fully visible’ state. There were significant differences among observers, which may be due to observer-specific variation in search technique -- the observers varied in how well they achieved the protocol guidance.

## Introduction

The Southeastern United States is a global hotspot for Chelonian diversity, ranking second only to Southeast Asia in species richness (Mittermeier et al. 2015). However, richness could possibly decline by up to 38% by 2080 (Ihlow et al. 2011). Eastern box turtles (*Terrapene carolina carolina*) are native to the Eastern United States, and like other turtle species, face threats from human development, road collisions, and illegal collection (Converse et al. 2005; Nazdrowicz et al. 2008; Currylow et al. 2011). According to the International Union for Conservation of Nature (IUCN), populations of all species of box turtles have declined 30% in the previous 100 years (van Dijk, 2011). Their longevity, late maturation, and high juvenile mortality exacerbate the effects of newly introduced stressors from the human environment (Erb and Roberts 2023). At U.S. Fish and Wildlife Service (USFWS), Patuxent Research Refuge in Central Maryland, Eastern box turtle population densities have dropped from 4 to 5 turtles per acre in the 1940s to about 1.2 turtles per ha (0.55 turtles per acre) today (Royle and Turner 2022). Declines at the Patuxent Research Refuge are consistent with range wide declines of the species (Hall et al. 1999; Kemp et al. 2022). As a result, there has been increased attention focused on developing monitoring efforts for Eastern box turtle populations (Roberts and Erb 2023).

Collecting data in a statistically rigorous manner is the first step towards understanding population status and achieving meaningful conservation of threatened species. For Chelonians, population data are often collected using visual search surveys (Willson and Dodd 2016), and the resulting data are analyzed in one of two ways: 1) Capture-recapture (CR) and related models (such as distance sampling), to estimate population size or density (Converse et al. 2005; Royle and Turner 2022), or 2) occupancy models, to estimate distribution and factors that affect probability of species occurrence (Langtimm et al. 1996; Erb et al. 2015). Probability of detection is a central concept in both classes of methods, and it is important to understand how detection probability varies over space, time, and habitat (Refsnider et al. 2011). Studies of reptiles and amphibians are particularly subject to detection bias and undercounting (Mazerolle et al. 2007). While visual surveys and occupancy methods remain popular with mammals, both methods may miss small and rare species that tend to have lower encounter rates, like many reptiles and amphibians. Mazerolle et al. (2007) found that methods involving marking or resighting individuals underestimated population metrics when failing to account for detection probability. In addition to the nature of herpetofauna, observer bias, habitat, and seasonality may also skew detection rates (Anderson et al. 2001; Gardner et al. 1999; Kéry et al. 2011).

A useful conceptual model of detectability in animal populations is the availability/perception conceptual model (Marsh and Sinclair 1989; Kendall et al. 1997, Bailey et al. 2004, Diefenbach et al. 2007) under which it is recognized that detection bias arises from two distinct processes: availability and perception. The availability (or its complement, temporary emigration) process determines when individuals in the population are available for detection by observers because of conspicuous behaviors (such as basking or foraging) or, conversely, are unavailable for detection because they are engaging in unsuitable behaviors such as brumation. The second process, perception, is the visual detection of animals by the observer given that they are available to be detected. In aerial surveys of waterfowl, Marsh and Sinclair (1989) defined perception bias as “the proportion of groups of the target species that are visible in the transect yet missed.” In the context of aerial surveys (and terrestrial turtle surveys), an animal being available for detection is one that is physically conspicuous to the observer. Such individuals may therefore be detected, or not, by the observer. Animals that are not available by virtue of being underground or buried in leaf litter cannot be detected by observers in ordinary visual search surveys. The precise interpretation of ‘available’ differs between search methods (e.g., humans vs. dogs). For example, in studies of salamanders (Bailey et al. 2004), individuals are inconspicuous on the ground, but become detectable once a cover object is lifted. It is important to understand both components of detectability because availability is driven largely by biological processes related to behavior and environmental conditions, whereas perception is driven more by factors related to observer and survey method.

One way to investigate detection probability experimentally is to simulate live population survey techniques with a controllable or measurable proxy (Fuentes et al. 2015). Two numbers are required to measure detection probability: A count of detections, via a systematic search method, and a known pool of available objects from which detections are made. Although various detection methods exist, simulating a known pool for an open system has proven more difficult. Previous studies reported low Chelonian detection probabilities using radio tagged live turtles (Refsnider et al. 2011) and others have corrected inflated population estimates using human-constructed model turtles to quantify human perception bias (Fuentes et al. 2015).

We initiated a field experiment to evaluate perception bias in visual search surveys of Eastern box turtles (hereafter “box turtles”). We surveyed plots with randomly placed museum collection box turtle shells in visible and partially visible states. Use of museum shells simulate a closed population where individual behavior is known and controlled. Our experience with surveys of live free-ranging Eastern box turtle is that most detected turtles are conspicuous on the forest floor, although we expect some conspicuous turtles are not detected. However, partially visible turtles in above-ground cover or mostly buried in leaf litter appear to be detected at a much lower frequency. Here, we attempt to determine if there are significant causes for variation in detection of box turtles during visual search-encounter surveys, and quantify the percentage of conspicuous box turtles that may go undetected by observers. We expect that individuals that occupy a study site may inhabit one of three states, consistent with the availability/perception model of detection probability: They may be completely invisible to observers, and undetectable without invasive searching (moving leaf litter, debris, etc.); They may be conspicuous and observable, on the surface of the leaf litter, foraging, basking or moving about their home ranges; Or third, there may be an intermediate state which is partially observable, comprising individuals that have some portion of their bodies exposed and thus susceptible to individual detection. We predict that partially observable box turtle shells (hereafter “shells”) should be detected at a lower rate than fully visible shells, and we hypothesize that experienced observers should be similar in their abilities to detect box turtles. We expect that any differences among observers should be explainable by measurable attributes of their search method, which we attempted to characterize using GPS search track data.

## Methods

### Plot Description

Detection surveys occurred in three plots delineated by orange flagging (Fig. 1). All plots were adjacent to powerline clearings on the U.S. Fish and Wildlife Service’s Patuxent Research Refuge in Laurel, Maryland. The refuge is located in a suburban area between Baltimore, Maryland, and Washington, D.C. (approximate refuge centroid 39.064N, 76.784W). Eastern box turtle monitoring has occurred on the refuge since the 1940s (Stickel 1950).

**Figure 1.**
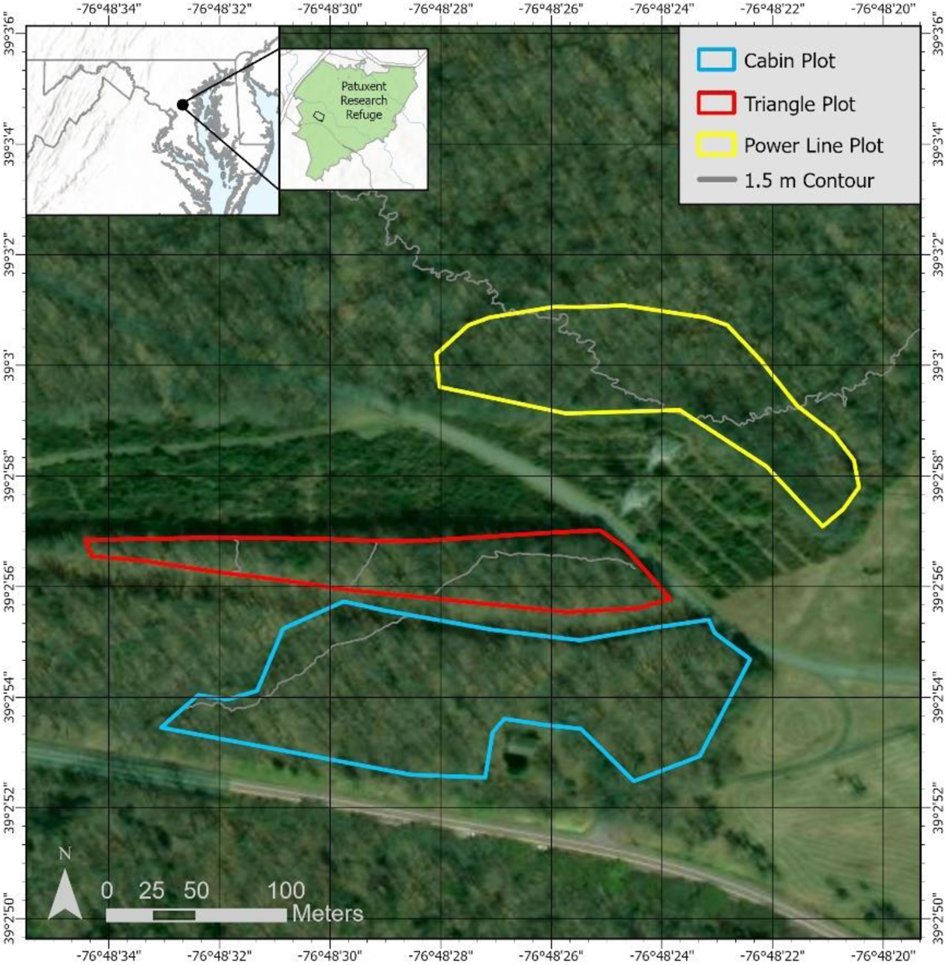
Map of Survey Plots on the U.S. Fish and Wildlife Service, Patuxent Research Refuge, Laurel Maryland, USA.

Two plots (“Triangle” and “Cabin”, see Fig. 2) were long and narrow totaling approximately 0.69 and 1.56 hectares, respectively. Both consisted of American beech (*Fagus grandifolia*) dominated upland with minimal understory, limited ground cover, fallen logs, and a small seasonal stream flowing through both. The third plot (“Powerline”) was rectangular totaling 0.94 hectares and was adjacent to a seasonal wetland. Beeches and American sweetgum (*Liquidambar styraciflua*) predominated, with minimal understory but with some dense patches of ferns.

**Figure 2.**
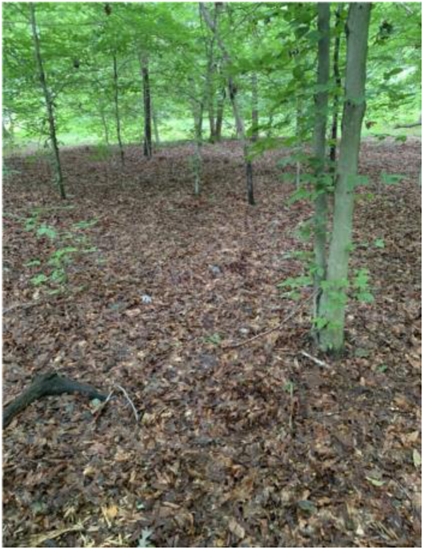
Example of the upland beech (*Fagus grandifolia*) habitat present in the Cabin survey plot in the U.S. Fish and Wildlife Service, Patuxent Research Refuge, Laurel, Maryland, USA. See Figure 1 for mapped location of the Cabin survey plot. *Photo: E. Wapman*.

### Shell Placement

Eastern box turtle museum collection shells were obtained from Towson University (Towson, MD) and Jug Bay Wetlands Sanctuary (Lothian, MD). The shells, varying in size and color, were previously shellacked (i.e., coated in a clear adhesive) for preservation. We do not believe this significantly affected their detectability, although of some effect may be possible. The number of shells placed varied each week and varied by plot, with a minimum of 3 shells and a maximum of 13 in any one plot during a survey event (defined here as the search of a single plot by a single observer). Typically, the shells were deployed to the 3 study plots on a weekend or Monday, and remained for the week so that each observer could survey each plot during the week at their convenience. The number and locations of shells were pseudo-randomly generated weekly by A. Royle with effort to minimize his knowledge about shell placement. Specifically, points were simulated and plotted using R (R Development Core Team 2019), and each map was viewed for only a few seconds before it was pasted into a document for the 1st and 2nd authors to distribute shells. In total, 116 shells were placed within 3 plots across the 5 weeks (Table 1) between June 13th and July 15^th^ 2022. The distribution of shells slightly favored the partially visible state (Table 2). Shells were randomly chosen every week to ensure different ones were used in each plot. Shells were placed in one of two states to imitate how live box turtles are encountered in population surveys based on our experience. “Visible” shells were placed on top of the forest floor, representing foraging, basking, or moving turtles. “Partially visible” shells were nestled against nearby logs or trees or buried in leaves covering the marginal scutes to a variable degree (Fig. 3).

**Figure 3.**
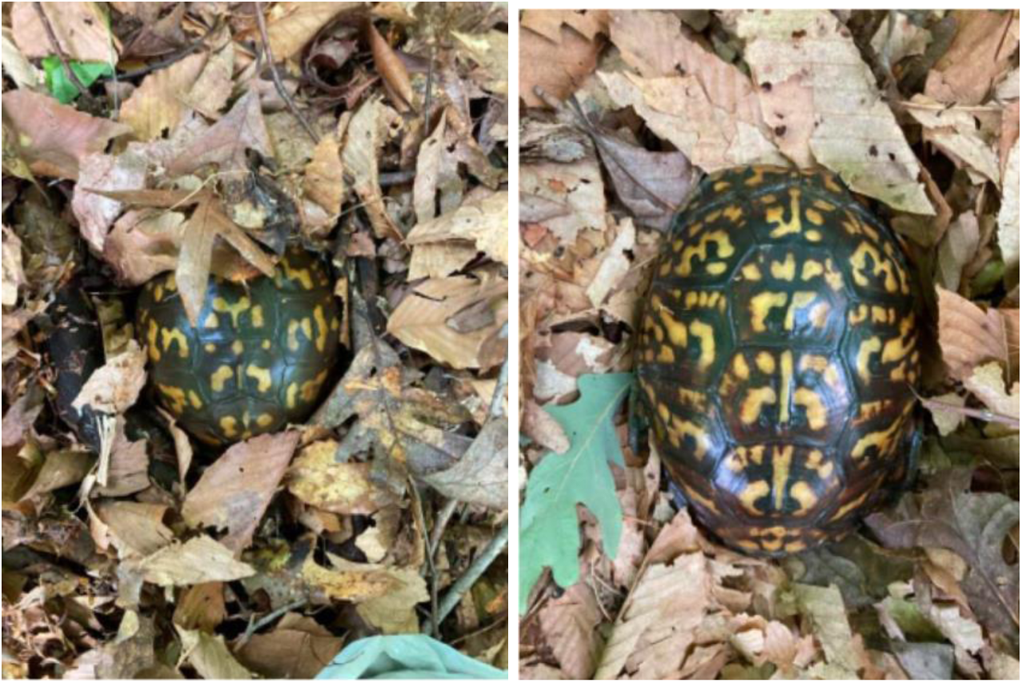
Example of a partially visible (left) vs fully visible (right) Eastern box turtle (*Terrapene carolina carolina*) shell placement for the experimental evaluation of box turtle detection in visual search surveys.

**Table 1.** Number of shells of Eastern box turtles (*Terrapene carolina carolina*) deployed per week in each of three survey plots between June 13^th^ and July 15^th^ 2022 in the U.S. Fish and Wildlife Service, Patuxent Research Refuge, Laurel, Maryland, USA. See Figure 2 for mapped location of the survey plots.

| Week | Cabin Plot | Power Line Plot | Triangle Plot | Total |
| --- | --- | --- | --- | --- |
| 1 | 8 | 9 | 3 | 20 |
| 2 | 8 | 7 | 4 | 19 |
| 3 | 10 | 9 | 3 | 22 |
| 4 | 10 | 11 | 4 | 25 |
| 5 | 13 | 12 | 5 | 30 |

**Table 2.** Eastern box turtle (*Terrapene carolina carolina*) shell ‘state’ by plot, total over the 5-week study period between June 13^th^ and July 15^th^ 2022 in each of three survey plots in the U.S. Fish and Wildlife Service, Patuxent Research Refuge, Laurel, Maryland, USA. See Figure 2 for mapped location of survey plots.

| Plot | Visible | Partially Visible |
| --- | --- | --- |
| Triangle | 22 | 27 |
| Powerline | 22 | 26 |
| Cabin | 8 | 11 |

### Shell Detection

Up to four biologists surveyed the three plots each week. Surveyors documented their search track with GPS (Gaia GPS) on their smartphone. Surveyors were instructed to achieve uniform coverage of the plot by envisioning a 5 m buffer on either side of their search track. There was no formal data-based reason for choosing this 5 m instruction other than for observers to have a consistent instruction to follow, although it was close to the mean detection distance (about 4.3 m) reported in Royle and Turner (2022). Detection almost surely depends on distance and thus no specific buffer will yield equal detection of box turtles. No search time constraints were imposed, consistent with our operational field protocol. When a surveyor detected a shell, they measured the distance between their location and the shell (not analyzed here), took photographs of the shell *in situ*, and recorded whether they determined it was visible or partially visible. Surveyors (except Royle) had no prior knowledge of the number of shells in each plot and did not survey plots where they had placed shells. All four surveyors were experienced box turtle searchers and have conducted operational surveys in these plots.

### Statistical Methods

We used observer search tracks to define covariates that might account for heterogeneity in detection probability of shells. Specifically, we placed a 5 x 5 m box around each of the deployed shells and used this to define covariates of searcher proximity to each shell. First, we computed ‘coverage’ -- the area of the 5 x 5 m box covered by a buffered track (Fig. 4). We initially used a 5 m buffered track, and thus the values of the coverage covariate ranged from 0 to 25 m^2^. However, coverage based on the 5 m buffered track produced values that were highly concentrated at the maximum value (25 m^2^) and thus were not an ideal covariate to measure search efficacy (i.e., did not capture variation among shells well). Thus, we computed a similar coverage covariate using a 2 m buffer which produced more variability in the covariate values to better represent the exposure of each shell to detection. This process is depicted in Fig. 4, which shows a single buffered track (2 m) for one of the observers during their search of the powerline plot. All geographic analysis calculations were done in the R package (R Development Core Team 2019).

**Figure 4:**
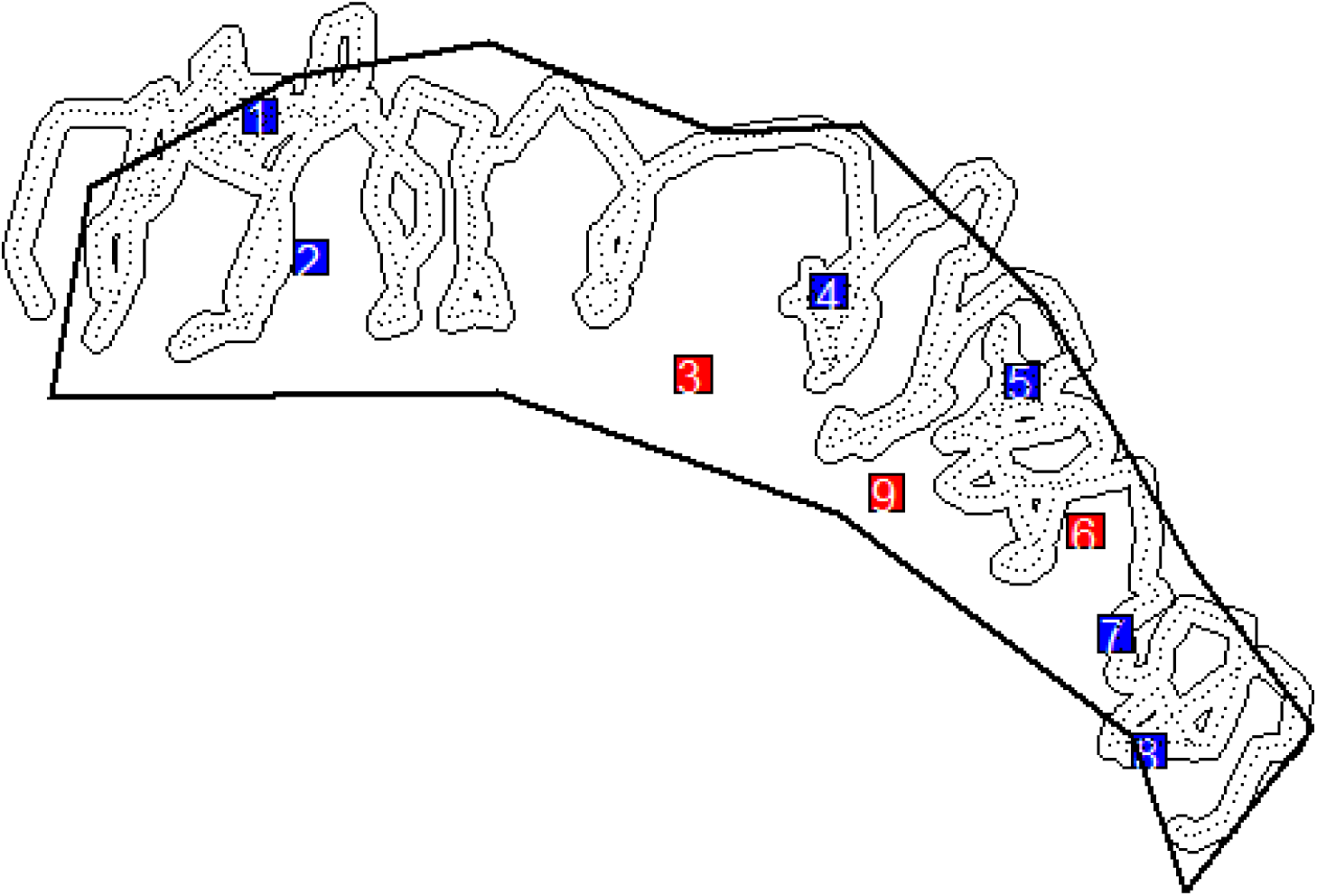
A search track with a 2 meter buffer for one of the searches of the Powerline survey plot between June 13^th^ and July 15^th^ 2022 in the U.S. Fish and Wildlife Service, Patuxent Research Refuge, Laurel, Maryland, USA. See Figure 2 for mapped location of survey plot. Detected Eastern box turtle (*Terrapene carolina carolina*) shells are shown in blue, shells that were not detected are shown in red. Bounding boxes around each shell are 5 x 5 m square polygons. See Figure 2 for mapped location of survey plot.

We created a second covariate, ‘points’, where we represented each search track by a set of regularly spaced points, using a constant intensity of 20 points per search minute. We tallied up the number of points in each 5 x 5 m shell grid box as a surrogate measure of composite time and effort spent in the vicinity of each shell. In the model fitting (see below), we used a square-root transformation of this variable due to extreme right skew.

To evaluate differences in search efficacy among observers, we also computed 3 plot-level covariates for each observer’s search as a summary of their overall search efficiency: the total proportion of plot area covered, computed by intersecting the 2- or 5-m buffered track with the plot boundary, and the total time spent surveying the plot.

We used logistic regression to model the probability of detecting shells. Let *y_ij_* denote the observed outcome (*y* = 1 if detected, *y* = 0 if not detected) for shell *i* = 1,2, …, 116 (total shell sets over the 5-week study) and observer *j* = 1,2,3,4. We assume that *y_ij_* is a Bernoulli outcome with parameter *p_ij_*, the probability that a shell is detected. Covariates are modeled as a linear function on the logit-transformation of *p* as follows:

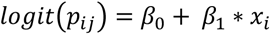

where *x_i_* is the value of some covariate for shell *i*. We considered the three covariates related to the observer search effort as defined above: ‘coverage’ is the area of the 5 x 5 m box around each shell that intersects the 2 m buffered search track, ‘points’ is the number of track points within the 5 x 5 m box (using a 20 point per minute density of regularly spaced points to define the GPS track), and ‘state’ which is the visible or partially visible state of each shell. The first two covariates were modeled as continuous covariates, the covariate ‘state’ was coded as a factor with the intercept of the model corresponding to the level ‘partially visible’. We also considered an observer effect (a factor having 4 levels), and we considered a plot effect (3 levels). We carried-out model selection in two stages. In the first stage we defined models that included various combinations of the experimental design variables: plot, observer, and state (7 models total, including the null model having no effects). The top two models from stage 1 were used as a basis for stage 2 model selection, including combinations of the shell-level covariates. A total of 13 logistic regression models were fit (see Results). Models were ranked using Akaike information criterion (AIC) score (with small-sample correction, AICc) to evaluate important effects and produce a ‘best’ model.

## Results

Four observers participated in at least 6 total plot searches (out of 15 plot-week sets). The number of shells each observer was exposed to was: 116, 60, 56, 55. The variation among observers is because observer 1 was always a searcher (all 5 weeks and 3 plots) whereas observers 2 and 3 alternated setting up shells each week, and observer 4 only participated in the last 2 weeks of the study as a trained observer. Overall, 49.8% of placed shells were detected by surveyors (143 out of 287). Visible shells were detected 21.5% more often than partially visible shells (63% for visible vs. 41.5% for partially visible, Table 3). Observers detected visible and partially visible shells in differing proportions. The largest difference between visible and partially visible shells was for Observer 1 who detected visible shells at a rate of 37.5% higher than partially visible shells, perhaps attributable to senescence and declining peripheral vision of that observer. The other observers had more similar detection rates among visible and partially visible shells. During the experimental surveys of the three plots, twelve live box turtles were incidentally encountered.

**Table 3:** Proportion of partially visible and (fully) visible Eastern box turtle (*Terrapene carolina carolina*) shells detected by each of four observers between June 13^th^ and July 15^th^ 2022 in the U.S. Fish and Wildlife Service, Patuxent Research Refuge, Laurel, Maryland, USA.

|  | Partially Visible | Visible |
| --- | --- | --- |
| Observer 1 | 29.6% | 67.1% |
| Observer 2 | 41.4% | 48.9% |
| Observer 3 | 49.5% | 78.3% |
| Observer 4 | 45.7% | 57.9% |
| **Total** | **41.5%** | **63.0%** |

A summary of the survey-specific covariates (total time and fraction of plot area covered using both the 2 m and 5 m GPS track buffer) for each observer are shown in Appendix Table 1. Total survey time ranged from 68 to 111 minutes for the Cabin plot (mean 88.7), 46 to 169 minutes for the Powerline plot (mean 73.2), and 40 to 87 minutes for the triangle plot (mean 61.3). Proportion of plot area covered using the 2 m buffered track ranged from 0.40 to 0.60 for the Cabin plot (mean 0.51), 0.39 to 0.68 for the Powerline plot (mean 0.58) and 0.42 to 0.63 for the Triangle plot (mean 0.52). For the 5 m buffered tracks the proportion of plot area covered ranged from 0.77 to 0.99 (mean 0.89) for the Cabin plot, 0.70 to 1.00 (mean 0.92) for the Powerline plot, and 0.70 to 0.95 (mean 0.86) for the Triangle plot. Thus, there was variation in observer effectiveness searching plots, with considerable variation both in search time and the fraction of the plot covered by the observer summarized in Table 4. For example, Observer 1 and Observer 2 achieved better coverage than Observers 3 and 4, on average. Observer 1 typically had the lowest time per unit area compared to the other 3 observers but covered more of the plot.

**Table 4.**
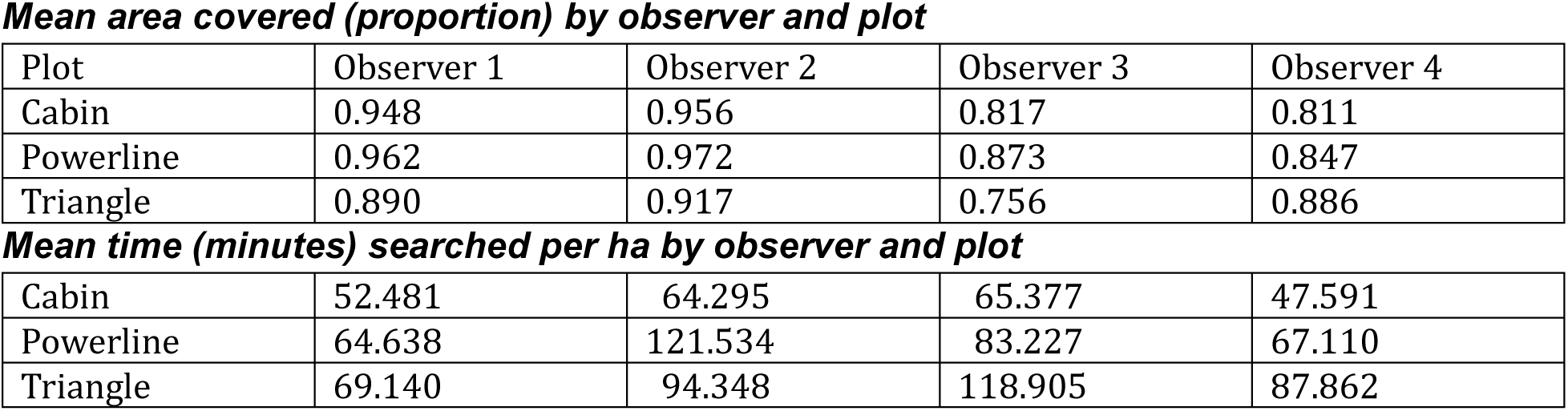
Observer summaries of proportion of area coverage (top portion) and time searching (minutes) per ha (bottom portion) for Eastern box turtle (*Terrapene carolina carolina*) shells on four study plots by four observers between June 13^th^ and July 15^th^ 2022 in the U.S. Fish and Wildlife Service, Patuxent Research Refuge, Laurel, Maryland, USA. See Figure 2 for mapped location of survey plots.

| Plot | Observer 1 | Observer 2 | Observer 3 | Observer 4 |
| --- | --- | --- | --- | --- |
| Cabin | 0.948 | 0.956 | 0.817 | 0.811 |
| Powerline | 0.962 | 0.972 | 0.873 | 0.847 |
| Triangle | 0.890 | 0.917 | 0.756 | 0.886 |
| ***Mean time (minutes) searched per ha by observer and plot*** |  |  |  |  |
| Cabin | 52.481 | 64.295 | 65.377 | 47.591 |
| Powerline | 64.638 | 121.534 | 83.227 | 67.110 |
| Triangle | 69.140 | 94.348 | 118.905 | 87.862 |

Logistic model definitions and AIC summaries are shown in Table 5. For phase 1 of the model fitting, the top model contains ‘state’ only, and the 2nd best model has state + observer. We used the top 2 models (m1 having state only and m5 having state + observer) as initial models for the 2nd stage of model fitting where we considered combinations of the two shell-level effort covariates (lower half of Table 5). This produced a top model (m11) which had state, ‘points’ (square-root transformed) and observer effects. The full model output including parameter estimates and standard errors (SEs) for the top 2 models (m11 and m7) is given in Appendix 1. The effect on detection probability of a shell being in the visible state was highly positive, 1.02 (SE = 0.26). The coefficient on the square-root of points was 0.43 (SE = 0.11). A graphical representation of the fitted response of detection probability to sqrt(points) is shown in Figure 5. The 2nd best model did not contain observer effects, and the coefficients of ‘state’ and ‘points’ changed little in that model relative to the top model (Appendix 1).

**Figure 5.**
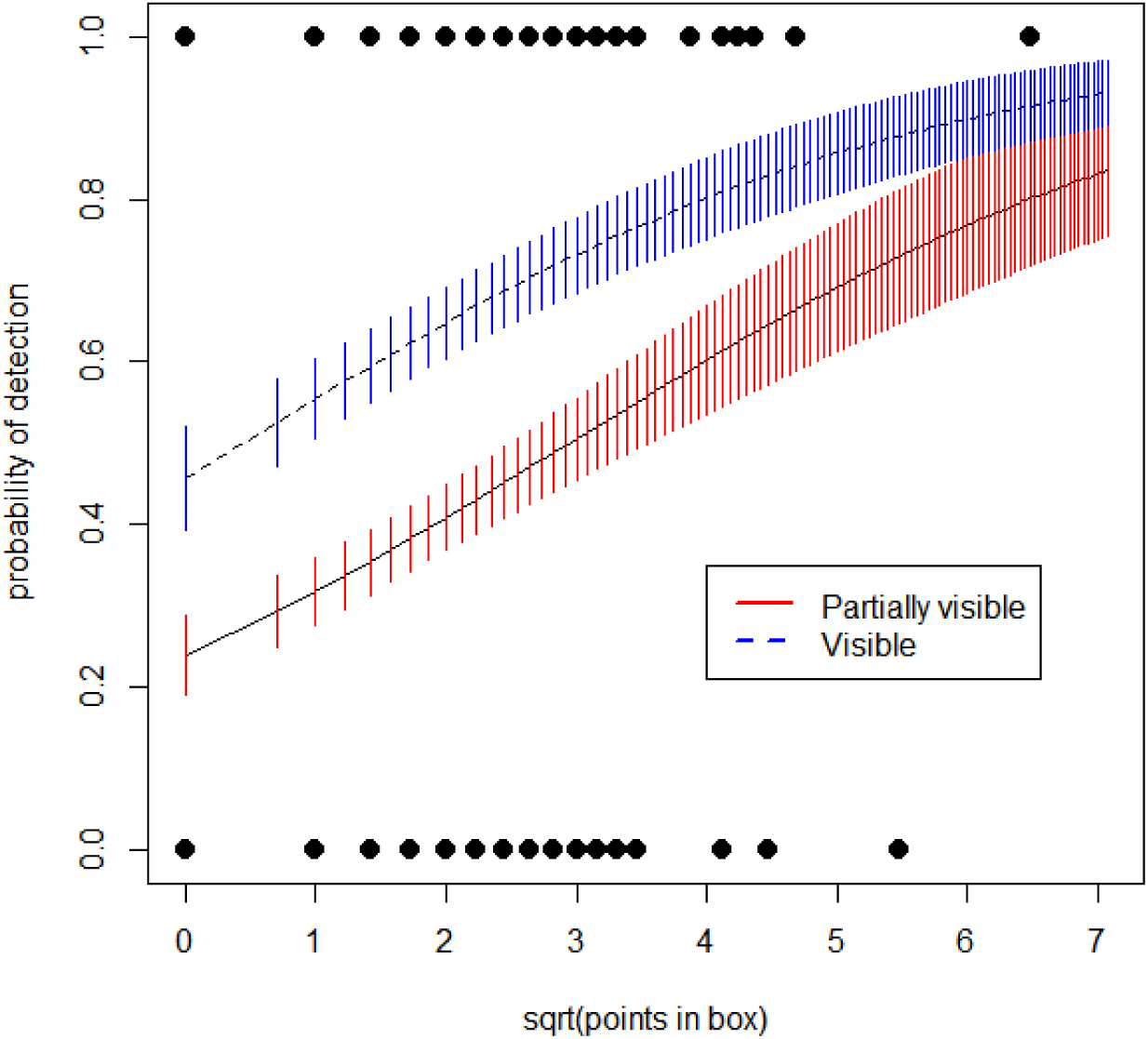
Fitted response curve for probability of detection of box turtle shells as a function of the search effort variable ‘points’. The ‘points’ variable represents the number (square-root transformed) of track points within a 5 x 5 m polygon surrounding each box turtle shell. Fitted response is shown for both visible and partially visible observation states. Solid points are the observed detection (y=1) or non-detection (y=0) events (multiple observations with the same value of the points variable pile up on top of each other). Vertical lines are predicted response +/- 1 SE.

**Table 5:**
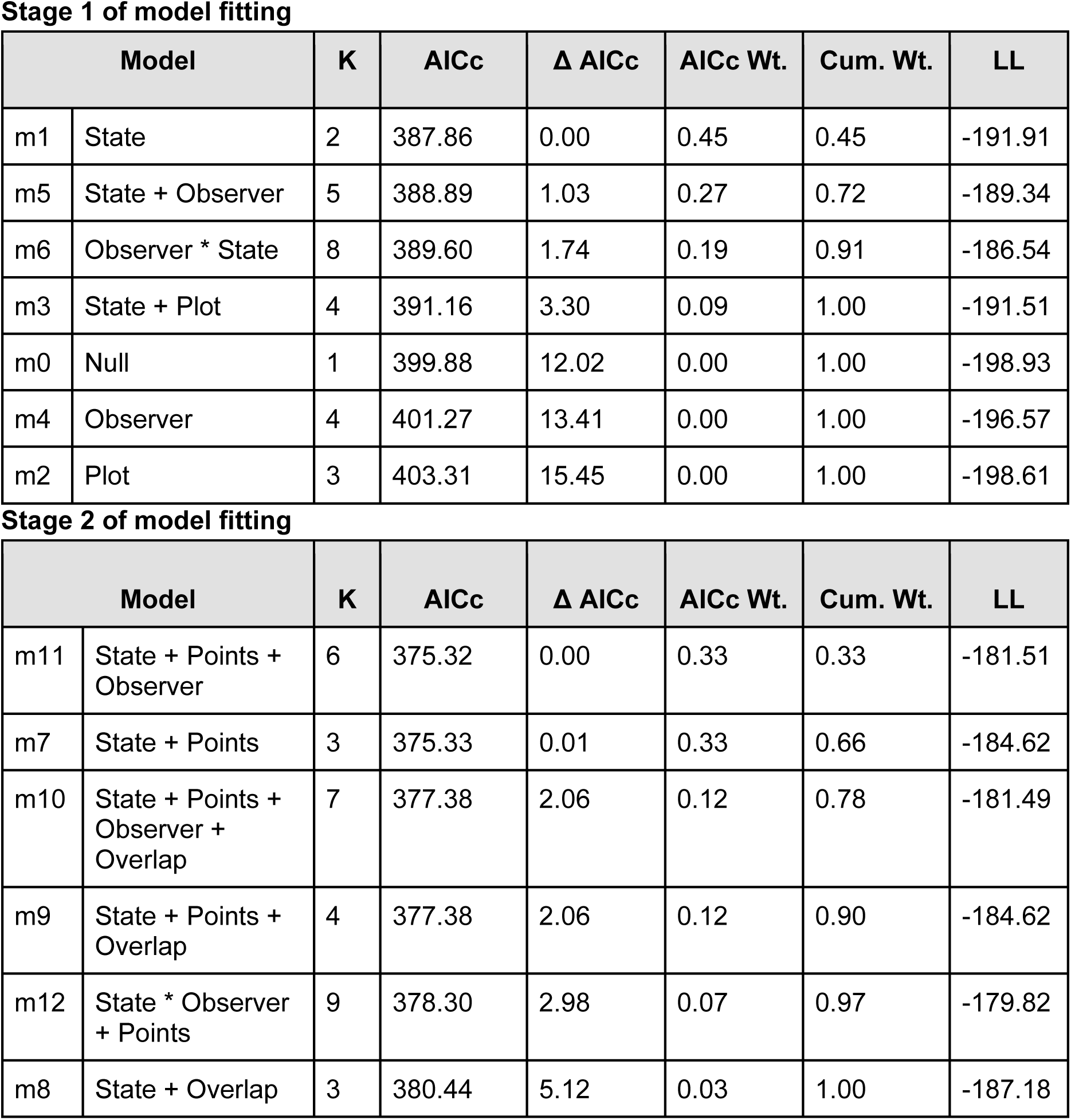
Logistic regression model selection results for Eastern box turtle detection study. Stage 1 considered models with the following effects on the probability of detecting a shell: Visibility state (partially and fully visible), observer (4 levels), and plot (3 levels). Stage 2 of the model fitting included covariates related to effort and search efficacy: ‘points’ is the number of GPS track points within a 5 x 5 m bounding box around each shell, and ‘coverage’ is the amount of the buffered GPS track that covers the bounding box. K is the number of parameters in the model, AICc is Akaike Information Criterion (AIC) score with small sample correction, AICc weights, cumulative weight, and log-likelihood are also given.

## Discussion

We implemented an experimental study to investigate the perception component of detection probability in visual search surveys of Eastern box turtle shells. Overall, about 50% of shells were detected across 4 observers (143 detections out of 287 trials), and this varied by visibility state (63% of visible shells, 41.5% for partially visible shells detected). This variation in detectability suggests that trained observers using our search protocol may miss about 37% of conspicuous box turtles in plain sight on leaf litter in a forest with predominantly open under-story. Furthermore, our results indicate there is variation among observers in the perception component of detection probability. Although trained, the 4 observers in our study varied in age and experience detecting box turtles, and we believe this may explain some of the variation in detection among observers. However, there was also clear variation among observers in how effectively they surveyed the plots according to our protocol of achieving uniform coverage subject to a 5 m transect width. Both coverage of the plot and time-per-unit area varied among observers, and many shells were not well exposed to detection by the observer, and thus not detected (i.e., some shells were beyond the 2 m buffered search track; Fig. 4).This demonstrates that ‘spatial coverage bias’ (i.e., incomplete spatial sampling) is an important component of perception, analogous to non-detection of distant objects in line-transect surveys.

Our results also suggest that observer variation in perception may be important even accounting for spatial coverage. That is, our top model favored observer effects in addition to the spatial variable, ‘points’ (the square-root of the number of GPS track points in the 5 x 5 m box around each shell). Care should be taken when comparing count based relative abundance indices that are not properly adjusted for observer variation.

Our observers missed a much larger proportion of ‘partially visible’ shells compared to fully visible shells. These partially visible shells were meant to mimic turtles emerging, or otherwise partially visible due to inactivity (i.e., resting or in-form rather than foraging, basking or mating). Presumably this partially visible state represents a relatively smaller fraction of turtles that are observed in the field, but we have not collected specific data on this during our operational surveys. However, we cannot determine from our study the fraction of the *population* that might be in the different availability states at any given time, either partially or fully visible. The probability of availability could be quantified with telemetry -- individuals could be located by observers and then the availability status of the individual recorded. Indeed, Refsnider et al. (2011) studied a population of ornate box turtles (*Terrapene ornata ornata*) with telemetry and estimated the probability of detection of available individuals (i.e., perception). Specifically, they refer to ‘detection probability’ which is the detection probability of available individuals, confirmed by telemetry. They estimated the perception probability to be 0.03. We are skeptical of that estimate given our own estimate, which is more than 15 times larger (about 50% averaged over visible and partially visible states). Moreover, consistent with our findings, two other studies (Gardner et al. 1999, Anderson et al. 2001; see below) estimated much higher estimates of the perception component of detection probability.

Besides Refsnider et al. (2011), to our knowledge there are two other studies that characterize the perception component of detection probability for tortoises. Gardner et al. (1999) used plaster molds of geometric tortoise (*Psammobates geometricus*) shells and had a group of observers simultaneously search plots using standardized transect searches. They achieved about 60% detection of tortoise shells in open habitat and 57% in denser habitat. Their individual searcher results were more variable, but no observer detected more than 50% of the models, similar to our overall detection rate of 50% (across partially and fully visible states). Detection probability in Gardner et al. (1999) varied by the size of the model and habitat denseness although the authors did not conduct formal model selection or hypothesis testing to gauge the statistical significance of these effects. In our study, there was not a wide variation in shell sizes, but we believe the variation in size was consistent with adult live turtles in the study area. Anderson et al. (2001) used styrofoam models to study the detection of desert tortoises (*Gopherus agassizii)* using line-transect survey protocols. Their sub-adult size models were only slightly larger than the adult sized box turtles in our study (7 inches vs about 5-6 inches for box turtles). The proportion of subadult tortoises detected ranged from 0.39 to 0.65 for 12 teams of 3 people searching a long transect. On average, 17% of models on or near (within 5 m) the centerline were not detected. Broadly speaking therefore, we believe the results of both Gardner et al. (1999) and Anderson et al. (2001) are consistent with our results demonstrating that a significant proportion of visible shells are missed by observers (or teams of observers), including shells that observers come into close proximity with.

We recognize some important limitations in our study. First, because our box turtle shells were museum specimens, most of them were treated with a coating of shellac to varying degrees. Overall, we did not feel that their appearance was much different from shells we observed in the field. However, we cannot be certain of this, and it is possible that the real detection frequency could be less than what we observed in the field experiment. We hope to evaluate this question using unmodified shells if a supply can be obtained. Second, as with Gardner et al. (1999) and Anderson et al. (2001), we conducted our detection study with a higher density of shells than the natural density of turtles in our study area, which is about 0.55 turtles/acre. Thus, our plots are expected to contain between 0-3 turtles on average. Data from our study from 1999-2022 indicate that 17 individuals have been captured in the Powerline plot since 2019, 7 in the Cabin plot (since 2020), and 12 in the Triangle plot (since 2020). An experimental design with natural densities would use only 1-3 shells per plot, thereby requiring many more weeks to obtain a similar number of detection events as we obtained under supernatural densities of box turtle shells. Therefore, our choice to use a higher density of shells in the study was pragmatic, but one might expect different detection rates under more realistic density settings. Moreover, as noted by Gardner et al. (1999), use of higher experimental densities “… allows for finer distinctions to be made in the experimental results.” Essentially, we expect higher statistical power to detect effects for the limited time we had to conduct the experiment. Finally, we only used two observable states, what we called visible and partially visible. We imagine that the unobservable state in which turtles are in-form or otherwise obscured, is the most common state for turtles to be in on any given day. For practical concerns we did not study fully obscured shells because during an experimental study the probability of observing a shell in this state should be close to zero. Although, we occasionally detect a small number (1-3) of fully obscured (i.e., what we would define as unobservable or ‘unavailable’) turtles every year by random chance (e.g., by stepping on them). The best way to study the availability of turtles (that is, the proportion of time they are not in the unobservable state) is with telemetry. With telemetry turtles can be detected nearly 100% of the time, and their availability, degree of conspicuousness, or some other measure of how available individuals are can be recorded. To our knowledge, this has not been done.

In summary, total proximity of an observer’s search track to the turtle shell as measured by our ‘points’ covariate was a key factor affecting probability of detecting a box turtle shell. Visibility state was also highly significant, but this variable is not controllable by surveyors. On the other hand, proximity is essentially a design feature that can be controlled -- observers can spend more time and do a better job systematically searching a plot, thereby increasing proximity for all possible shell locations and realizing a concomitant increase in detection probability (Fig. 5). But increases in detection probability cost time and money and, in practice, you can moderate time/effort and still estimate population parameters, like density, population size, or occupancy, in a manner that incorporates imperfect detection. As in many other wildlife monitoring systems, we found variation in detection probability among observers. However, this effect was not strongly significant. Nevertheless, investigators should consider accounting for observer effects in models whether for relative abundance based on count indices (e.g., Sauer and Link 2011) or capture-recapture to estimate population size, especially when multiple observers are used who may be highly variable in their skill levels and experience. When feasible, evaluating search protocols that use experimental trials may help understand the efficacy of a monitoring program, and provide insight into plausible models for wildlife status and trend assessments.

## Acknowledgements

We thank Steve Kimble and Towson University and Mike Quinlan and Jug Bay Wetlands Sanctuary for contributing box turtle shells. We are grateful to Haley Turner, Jackie Guzy, and Meg Lamont for providing reviews of a draft. We thank USGS CSFP Internship program that provided funding for E. Wapman, and Univ. Richmond Department of Biology who provided funding for W. Heinle and N. Beswick. We thank Sandy Spencer and U.S. FWS Patuxent Research Refuge for access to the refuge for box turtle research. Any use of trade, firm, or product names is for descriptive purposes only and does not imply endorsement by the U.S. Government.

## Appendix

*Disclaimer: Any use of trade, product, or firm names is for descriptive purposes only and does not imply endorsement by the U.S. Government*.

Summary results (R output) for the top 2 models fitted to the Eastern box turtle (*Terrapene carolina carolina*) detection data.

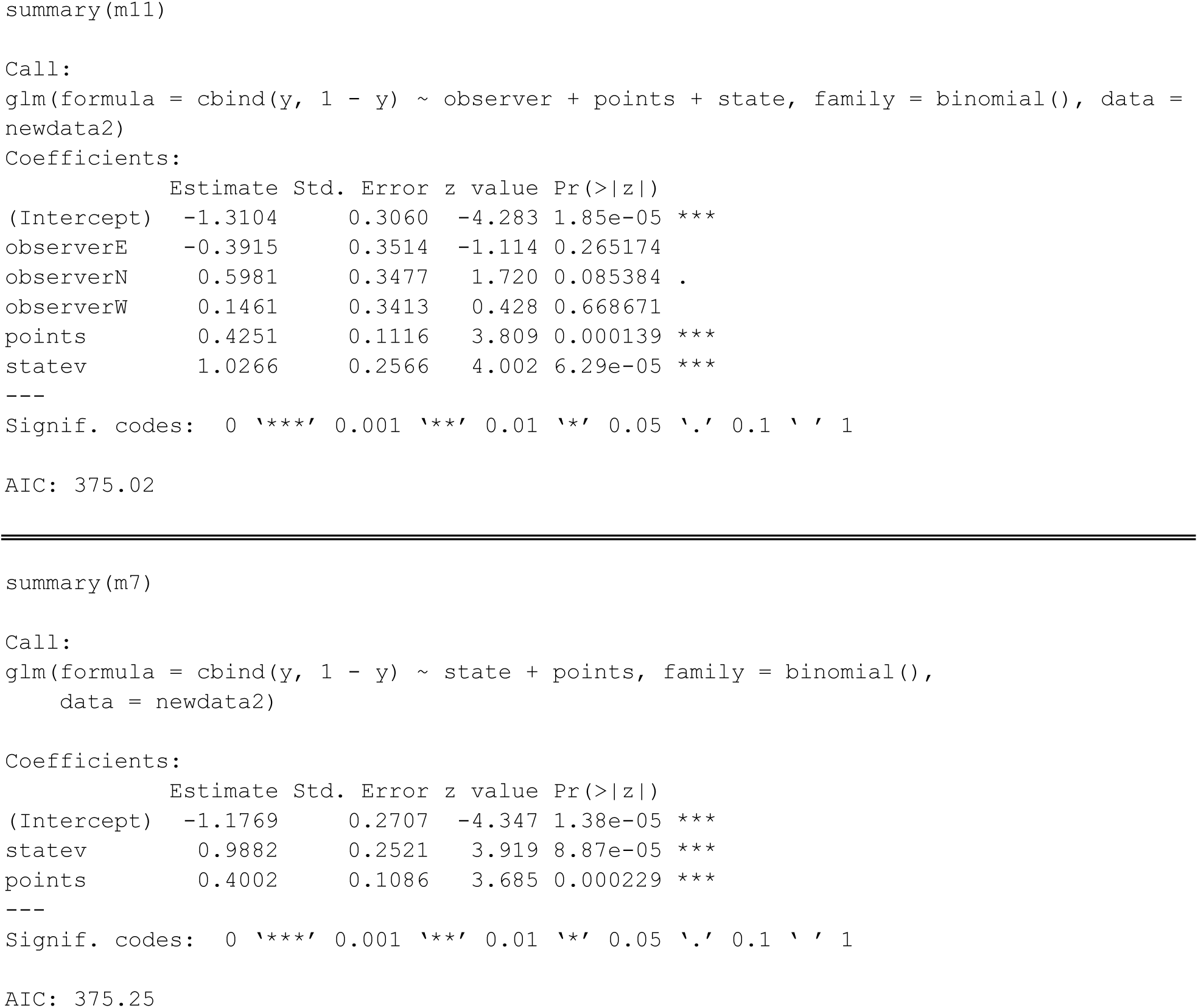

**Appendix Table 1:** Survey attributes for each observer’s search track. ‘plot’ values are Pow = Powerline, Cab = Cabin and Tri = Triangle plots as referenced in the main text; ‘observer’ references the individual conducting the survey (values 1-4); ‘survey’ is an integer representing the replicate sample of each plot (values 1-5); ‘totaltime’ is the time in minutes it took the observer to complete the search, ‘Areafrac2’ is the total plot coverage using a 2 m track buffer, ‘Areafrac5’ is the same using a 5 m buffer, ‘Area.cov2’ is the total area covered using the 2 m buffer and ‘Area.cov5’ is the total area covered using the 5 m buffer.

|  | plot | observer | survey | totaltime | Areafrac2 | Area.cov2 | Areafrac5 | Area.cov5 |
| --- | --- | --- | --- | --- | --- | --- | --- | --- |
| 1 | Pow | 1 | 1 | 59.0 | 0.563 | 4445 | 0.861 | 6796 |
| 2 | Pow | 4 | 1 | 46.6 | 0.230 | 1819 | 0.703 | 5551 |
| 3 | Pow | 1 | 2 | 46.0 | 0.764 | 6032 | 0.977 | 7717 |
| 4 | Pow | 3 | 2 | 79.1 | 0.630 | 4971 | 0.892 | 7040 |
| 5 | Pow | 1 | 3 | 55.1 | 0.734 | 5799 | 0.988 | 7804 |
| 6 | Pow | 4 | 3 | 57.3 | 0.704 | 5561 | 0.886 | 6997 |
| 7 | Pow | 1 | 4 | 80.8 | 0.867 | 6845 | 0.986 | 7784 |
| 8 | Pow | 2 | 4 | 168.7 | 0.797 | 6295 | 0.975 | 7698 |
| 9 | Pow | 3 | 4 | 77.4 | 0.658 | 5193 | 0.854 | 6741 |
| 10 | Pow | 1 | 5 | 62.9 | 0.944 | 7454 | 0.999 | 7888 |
| 11 | Pow | 2 | 5 | 59.8 | 0.842 | 6644 | 0.969 | 7651 |
| 12 | Pow | 4 | 5 | 85.3 | 0.180 | 1418 | 0.952 | 7518 |
| 13 | Cab | 1 | 1 | 74.0 | 0.794 | 12378 | 0.948 | 14785 |
| 14 | Cab | 3 | 1 | 107.8 | 0.702 | 10949 | 0.864 | 13482 |
| 15 | Cab | 1 | 2 | 84.6 | 0.812 | 12665 | 0.973 | 15183 |
| 16 | Cab | 4 | 2 | 68.3 | 0.544 | 8489 | 0.773 | 12053 |
| 17 | Cab | 1 | 3 | 86.5 | 0.792 | 12355 | 0.929 | 14489 |
| 18 | Cab | 3 | 3 | 87.1 | 0.571 | 8906 | 0.815 | 12712 |
| 19 | Cab | 1 | 4 | 83.9 | 0.792 | 12358 | 0.943 | 14709 |
| 20 | Cab | 2 | 4 | 109.0 | 0.762 | 11888 | 0.988 | 15406 |
| 21 | Cab | 4 | 4 | 80.2 | 0.626 | 9765 | 0.849 | 13251 |
| 22 | Cab | 1 | 5 | 80.4 | 0.637 | 9942 | 0.949 | 14800 |
| 23 | Cab | 2 | 5 | 91.6 | 0.820 | 12791 | 0.924 | 14410 |
| 24 | Cab | 3 | 5 | 111.1 | 0.490 | 7651 | 0.771 | 12025 |
| 25 | Tri | 1 | 1 | 43.9 | 0.608 | 4907 | 0.916 | 7387 |
| 26 | Tri | 3 | 1 | 87.3 | 0.444 | 3584 | 0.697 | 5624 |
| 27 | Tri | 1 | 2 | 64.0 | 0.694 | 5595 | 0.885 | 7139 |
| 28 | Tri | 4 | 2 | 46.5 | 0.595 | 4802 | 0.836 | 6746 |
| 29 | Tri | 1 | 3 | 41.2 | 0.591 | 4766 | 0.876 | 7061 |
| 30 | Tri | 3 | 3 | 87.4 | 0.535 | 4316 | 0.818 | 6596 |
| 31 | Tri | 1 | 4 | 49.7 | 0.739 | 5963 | 0.903 | 7284 |
| 32 | Tri | 2 | 4 | 71.6 | 0.701 | 5656 | 0.881 | 7104 |
| 33 | Tri | 4 | 4 | 74.8 | 0.729 | 5879 | 0.936 | 7546 |
| 34 | Tri | 1 | 5 | 39.6 | 0.494 | 3983 | 0.872 | 7033 |
| 35 | Tri | 2 | 5 | 58.6 | 0.825 | 6652 | 0.953 | 7689 |
| 36 | Tri | 3 | 5 | 71.4 | 0.514 | 4148 | 0.754 | 6079 |

